# SAKit: an all-in-one analysis pipeline for identifying novel protein caused by variant events at genomic and transcriptic level

**DOI:** 10.1101/2023.03.06.531407

**Authors:** Yan Li, Boran Wang, Zengding Wu, Shi Xu, Fenglei Cui, Caiyi Fei, Qiang Sun

## Abstract

**Summary:** Genetic modifications that cause pivotal protein inactivation or abnormal activation may lead to cell signaling pathway change or even dysfunction, resulting in cancer and other diseases. In turn, dysfunction will further produce “novel proteins” that do not exist in the canonical human proteome. Identification of novel proteins is meaningful for identifying promising drug targets and developing new therapies. In recent years, several tools have been developed for identifying DNA or RNA variants with the extensive application of nucleotide sequencing technology. However, these tools mainly focus on point mutation and have limited performance in identifying large-scale variants as well as the integration of mutations. Here we developed a hybrid <u>S</u>equencing <u>A</u>nalysis bioinformatic pipeline by integrating all relevant detection <u>Kit</u>s(SAKit): this pipeline fully integrates all variants at the genomic and transcriptomic level that may lead to the production of novel proteins defined as proteins with novel sequences compare to all reference sequences by comprehensively analyzing the long and short reads. The analysis results of SAKit demonstrate that large-scale mutations have more contribution to the production of novel proteins than point mutations, and long-read sequencing has more advantages in large-scale mutation detection.

**Availability and implementation:** SAKit is freely available on docker image (https://hub.docker.com/repository/docker/therarna/sakit), which is mainly implemented within a Snakemake framework in Python language.

## 1 Introduction

Numerous basic science questions regarding diverse human diseases, such as cancer, are related to novel proteins caused by base substitutions, deletion, insertion, frameshift, intron retention and alternative splicing, as well as novel unannotated open reading frames (nuORFs) translation at genetic and transcript level[1,2]. Because novel proteins are not included in the canonical human genome and not expressed in normal cells, they would not have immune tolerance, and could be easily recognized as foreign antigens. Novel proteins could be digested into short peptides, which are represented as T cell antigens by the host system, leading to consequential stimulation of cellular immune response and eradication of the cells expressing novel proteins [3]. Based on this principle, neoantigen-based immunotherapy has been developed by pharmaceutical companies or research institutions as a promising cancer treatment [4–6]. Additionally, novel proteins are also associated with other complex diseases, for instance, Alzheimer’s[7], type II diabetes[8], liver disease[9–14], kidney disease[15–20], Obesity[21–25], and cardiovascular disease[26]. Therefore, effective identification of novel proteins is imperative for understanding human diseases and developing corresponding therapies.

The traditional methods of identifying and characterizing protein are the combination of tandem mass spectrometry (MS) with subsequent database searching. However, if MS relies on reference databases containing regular sequences in the canonical proteome, it would be almost impossible to identify proteins outside the canonical proteome, especially those produced by patient-specific individualized mutants or variants. Moreover, by benchmarking with known protein repertoire, Lloyd D.Fricker[9] proposed that MS-based peptidomic approaches have other technical shortcomings: significantly lower detection rate in Cys-riched peptides, low detection sensitivity for peptides at low molecular weight (<500 Da) or high molecular weight (>3kDa). Obviously, the bias derived from technical issues limits the application of MS in identifying novel proteins. Therefore, new bioinformatic pipelines would be necessary for identifying novel proteins for immunotherapy.

In the past decade, the unprecedented improvement of Next-Generation Sequencing (NGS) platforms has paved the way to affordable sample-specific whole genome sequencing or total RNA sequencing. Undoubtedly, the advances and widespread application of NGS have accelerated revealing tremendous potential complexity of human proteome. In 2012, Wang et al.[10] proposed a protein identification workflow by constructing hypothetical protein sequence reference databases derived from RNA-seq data to facilitate deciphering mass spectrometry (MS) data, and the results demonstrated that such sample-specific reference databases could significantly increase peptide identification sensitivity, reduce protein assembly ambiguity, and further enable the detection of novel peptide caused by highly variant regions. Furthermore, this NGS-driven proteogenomic strategy has been increasingly applied to characterize protein repertoire in basic research or disease biology [11,12]. However, despite the improvements listed above, this method combining NGS and MS still suffers from several deficiencies:

1. NGS has inherent technical defects in detecting large structural chromosomal variations and transcriptic alternative splicing, due to the short read-length (generally no more than 150bp);
2. The identification protocol is redundant for most canonical proteins, most of which are required to be detected by both MS at protein level and NGS at RNA level. Although such redundancy benefits the characterization of smaller proteins[13], excessive sample input is mandatory to meet two detection protocol. But such sample input may not be constantly satisfied in all studies, thus it seriously limits the translational medical research from basic science to clinical application. Especially, in the situation that regenerative clinical samples are extremely precious, even the failure of processing a single sample may bring unacceptable consequences;
3. In terms of the nucleotide sequencing-based bioinformatic analysis workflow, most of them only analyze conventional transcripts in general or mainly focus on single or few variation categories; in other words, by design principle they cannot cover all sample-specific variations. Besides the limitation of short read-lengths in NGS platform, several other factors make it challenging to develop an analysis workflow covering all types of mutations relevant to novel proteins. We would explain these factors below:

a. Various categories of variants could all contribute to the production of novel proteins,e.g., base substitution, frame-shift, chromosomal instability, etc. The detection of each category of variants requires establishing a suitable analysis workflow customized for specific detection platform purposes. These tools are usually interdependent or intertwined, thus a complete analysis process involving distinct analysis on various variants would be invariably time-consuming and labor-intensive;
b. Calling and filtering that cover all variants often require massive computing power, especially when multiple samples are analyzed in parallel, and managing such time-consuming analysis steps is also a challenge;
c. The calling results of each variant category are extremely redundant and glutted with noise, implying high proportions of false positive results. Therefore, further fine filtering with proper parameters would be necessary for achieving accurate enrichment of true positive results.

Fortunately, the HIFI sequencing technology of the third-generation sequencer Pacbio platform is compatible with the characteristics of ultra-long read length and high accuracy. Therefore, it is especially suitable for detecting large fragment variants, alternative splicing isoforms, new open reading frames, etc. As long as sufficient sequencing depth of HIFI sequencing is guaranteed, point mutations can also be precisely detected. However, signals of point mutations are easily interfered with by random errors at a small length span. Therefore, if applicable, it would be ideal to combine NGS technology to obtain high-depth sequencing data to improve the detection accuracy of HIFI sequencing.

To break through the known limitations on protein identification, we developed a bioinformatics workflow named SAKit, which combines third-generation sequencing (HIFI) and second-generation sequencing (NGS) to generate a sample-specific hypothetical proteome. Besides an accurate analysis pipeline of the canonical transcriptome, SAKit also fully covers various alternatively spliced isoforms or variants that may generate novel proteins., A detection module is correspondingly developed for each specific variant category, and the results of each module are hierarchically filtered. Finally, the sequences comprising the hypothetical proteome derived from RNA sequencing are converted into fasta format. This process has been benchmarked against reference standards and showed high sensitivity and specificity. In general, we have developed an automated protein identification pipeline that is simple to use and excellent in performance.

## 2 Materials and methods

In this section, the framework and logic of the SAKit are summarized, as well as the dependence among different modules in the pipeline. Then we focus on the implementation of hierarchical filtering strategies in different modules to remove false hits. Finally, the methodology for benchmarking the key module and the entire pipeline is presented in detail.

### 2.1 Pipeline overview

Snakemake, a workflow management system, has several significant advantages: excellent memory management,high portability,high modularity,stable reproducibility. Therefore, SAKit uses the Snakemake as the management system, enabling the well-controlled and scalable pipeline execution in a high-performance computing environment. In terms of input data types, SAKit is compatible with the data of NGS (optional) and HIFI Sequencing; in terms of analysis procedures, SAKit includes multiple steps, e,g, quality control, calling, filtering, and annotation; in terms of variant categories, SAKit covers point mutation, insertion, deletion, frameshift, gene fusion, retained intron and alternative splicing. All these variant categories contribute to novel transcripts or proteins, which we will hereinafter refer to as Novel Variant Isoform (NVI); in terms of tool kits, multiple open-source software tools have been leveraged in the pipeline.

The SAKit pipeline takes a subreads.bam file of the raw data generated by the HIFI sequencing as input, together with a configuration file containing various optional parameters and necessary path of reference and annotation files, as well as raw data generated by NGS platform if available. For the first time locally deploying the pipeline before the test run, users may either directly set the parameters to the default values or modify the values for actual situations. After local deployment is completed, in each subsequent analysis, users only need to specify the path of sample-specific raw data to run the analysis.

The SAKit pipeline has more than two dozen rules, which can be divided into four parts:

1. quality control and pre-processing of raw data
2. calling of novel variant isoform (NVI)
3. hierarchical filtering and annotation for calling results
4. translating discovered transcripts to protein sequences in fasta format and identifying novel proteins outside canonical human proteome.

Open-source software provided by PacBio is applied in the pre-processing of part 1). We employ CCS [14] to obtain high-accuracy consensus HIFI reads from the original tandemly repeated subreads. Typically 99.9% accuracy can be obtained from the subreads with more than 3 circles after consensus calling. Actually, CCS analysis is the most time-consuming step in the whole pipeline. Then lima[15] is applied to trim barcodes so that multiple samples constructed in a single library could be distinguished, whereas this step can be omitted if barcodes are not employed when a library consists of only a single sample. Thereafter the refine module of isoseq3[16] software is applied to trim polyA tails and adapter sequences wrapped on both sides of HIFI reads to obtain full-length non-chimeric (FLNC) sequences; moreover, residual concatemers are also removed in this step. After obtaining the FLNC, the cluster module of isoseq3 is further used to cluster FLNC reads using hierarchical n*log(n) alignment and iterative cluster merging. In the novel variant isoform (NVI) calling step following pre-processing step described above, we utilized the *collapse_isoforms_by_sam.py* of cDNAcupcake [17], which takes bam file as input obtained by mapping cluster.fasta files to the pre-defined reference using minimap2[18], to collapse two or more identical transcript isoforms into a unique transcript isoform.[Fig2A] Simultaneously, the number of full-length reads corresponding to different unique isoforms was also obtained, which are important supporting evidence for the called isoforms, and will be used as the parameter in the following filter step. On the other hand, gene fusions are considered to be another important source of novel proteins or chimeric protein[19], and was called by utilizing *fusion_finder.py* of cDNA_cupcake.

**Figure 1.**
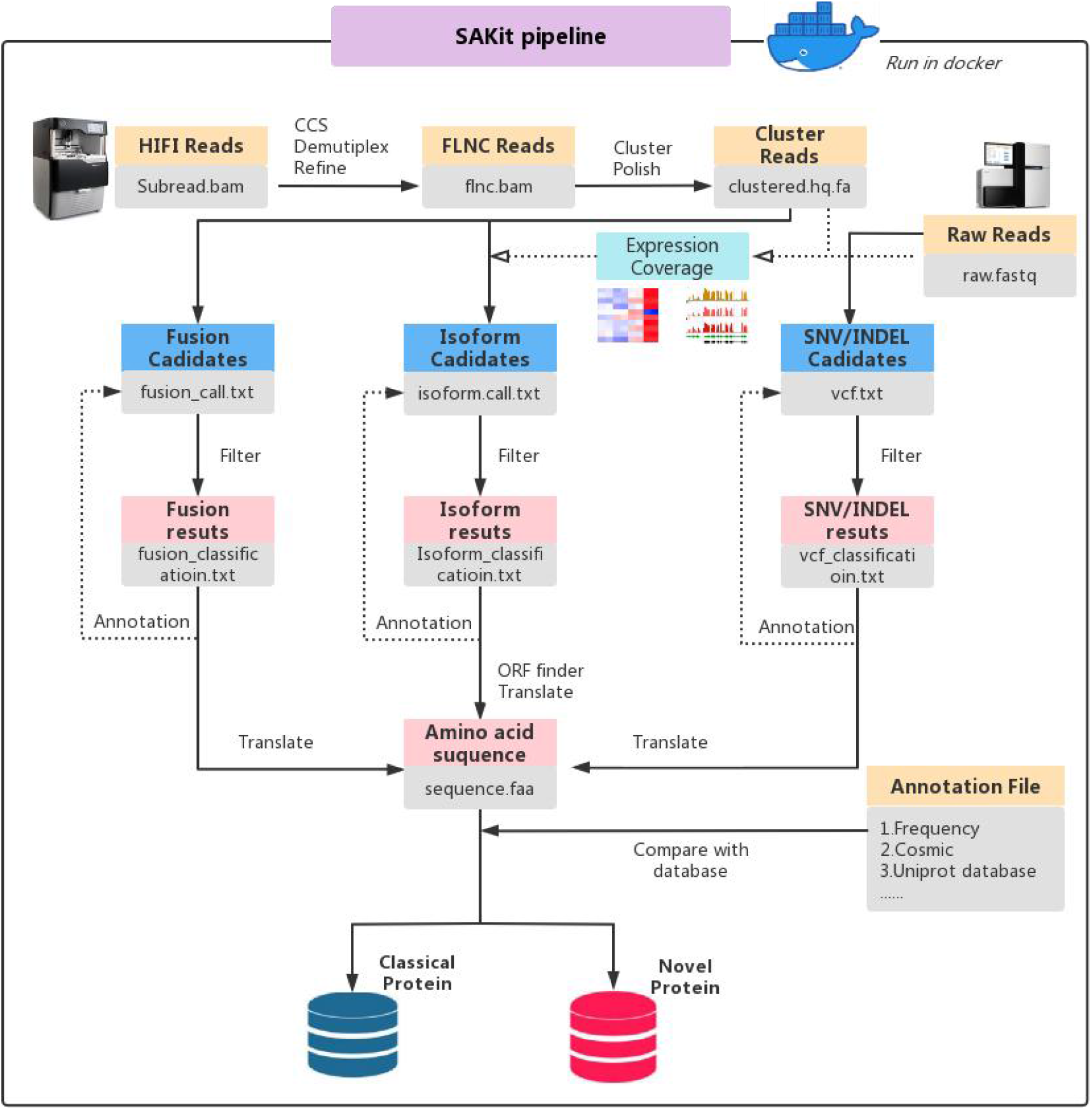
Workflow of SAKit for all-in-one calling all novel variants and isoforms containing single nucleotide variation(SNV), Insertion–deletion mutations(INDELs), genes or transcripts fusion(FUSION), ISOFORM of intron-retention and alternative splicing simultaneously by comprehensive analysis of NGS and HIFI sequencing data. The intermediate steps of the analysis includes multiple strict filtration, and finally the result of low background noise is obtained. Finally, the converted amino acid sequences were compared with the protein database to select Novel or Classic protein.

**Figure 2.**
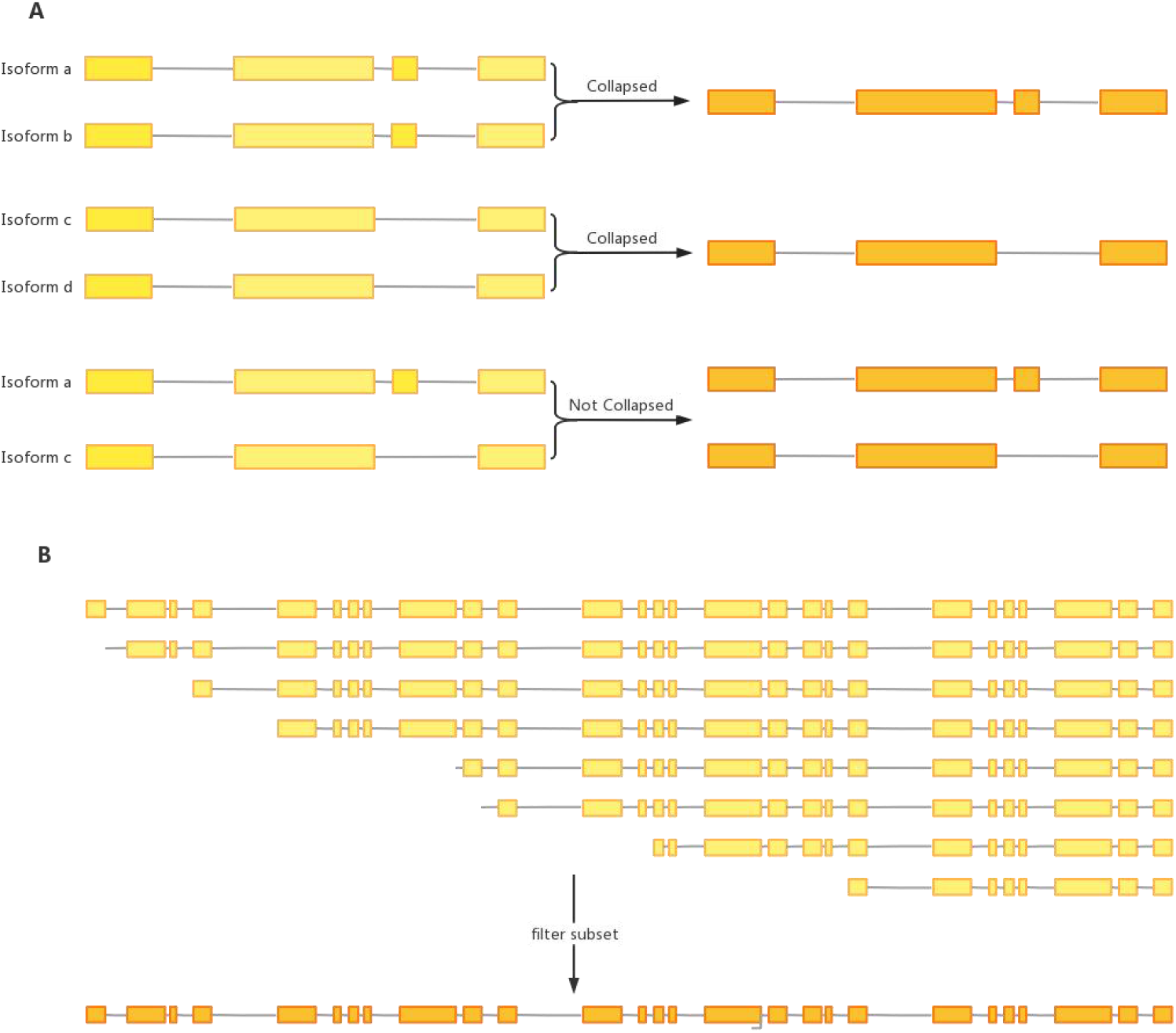
A) Shows the process of collapsing the long read sequence. Because sequence a and b, sequence c and d have exactly the same introns, they respectively collapse to get the only potential transcript candidates, while sequence a and c cannot further collapse because of their differences in introns. B) Shows that many sequences cannot be aligned at the 5 ‘end, but are neat at the 3’ end. Such sequences are biologically likely to be of the same original transcript with the 5 ‘degradation, so they are collapsed into a unique transcript isoform.

Nonsynonymous point mutations may induce immunogenicity: a peptide with a single amino acid difference from wildtype protein could be used as a neoantigen epitope in cancer vaccines[20–22]. Therefore, the SAKit pipeline also identifies SNV/INDEL variants to fully cover all possible novel transcripts producing proteins. The alignment result file of the NGS data is the input of vardict[23], whose output is the vcf file containing all variant candidates.

After preliminary results for all types of NVIs are obtained separately from corresponding toolkits. Due to the complexity and background noise, further enriching the true positive results by hierarchical filtering is necessary. Finally, the enriched NVI results are translated into protein sequences and compared with Uniprot or an in-house protein database to get the final results of novel or canonical proteins.

### 2.2 Hierarchical filtering

#### 2.2.1 Filtering of isoform/fusion

Since FUSION is also regarded as a particular type of isoform, its filtering logic and steps are basically the same as isoform. There are four steps to filter out the called isoforms[Figure3].

**Figure 3.**
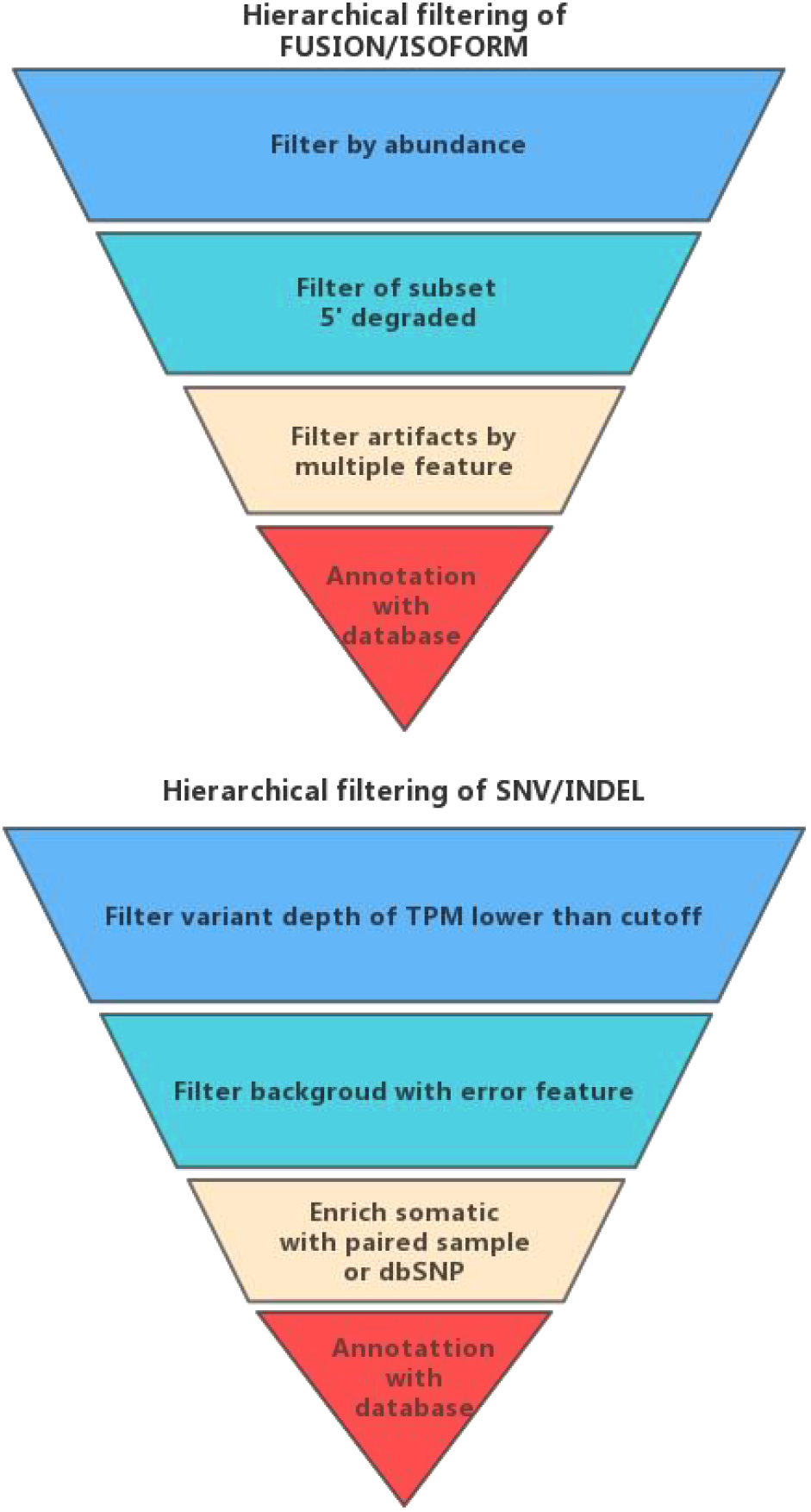
Hierarchical filtering for preliminary calling results of all novel variant isoform (NVI).The inverted triangle represents the logic of filtering. Each layer represents one step filtering operation. The values on the left and right sides are the representative number of results obtained by a clinical sample after each filtering step.

The first step is to deplete isoforms below the pre-set threshold (default 2) by abundance as a high-quality isoform should be supported by at least two FLNC reads[24]. Normally, the abundance of each called isoform could be directly quantified from PacBio HIFI sequencing data. Nonetheless if additional NGS sequencing data could be obtained from another aliquot of the same sample, it would provide an even more accurate quantification of isoforms’ abundance.

The second step is to deplete the subset of isoforms [Figure 2B] that have the same exons at 3’ end as the longest one but are missing some of the 5’ exons, and those 5’ truncated transcripts are likely due to degradation of the 5’ end of RNA during library construction[24]. Moreover, the more degraded the library, the greater the amount of transcripts filtered out due to 5’ truncation.

The third step is to deplete the “artificial” isoforms introduced by Chimerism in PCR, restriction enzyme ligation or sequencing error, etc. Such artificial isoforms are real signals called from the sequencing data, but they are noise to the samples and must be eradicated. Because a single feature could hardly identify random interference causing artificial isoforms, we use the machine learning-based SQANTI3 algorithm[25] to achieve this goal. SQANTI3 extracts up to 47 features including coverage, exon coordinate, supported reads number, etc., and effectively implements filtering based on these features.

The last step is the annotation with the database. This step can identify meaningful isoforms/fusion associated with clinical diseases, as well as potential therapeutic targets. The detail and accuracy of gene annotation results are highly dependent on the quality and quantity of the annotation database. For fusion gene annotation, we take the database of chimerDB4.0[26], which summarizes nearly 100k fusion candidates from patients in the TCGA project and including tens of thousands fusion genes with experimental evidences support. While appropriate isoform annotation databases are still lacking, although there are gradually some transcript isoform databases, they are quite incomplete and unfriendly to use. It is expected the emergence of high-quality transcript annotation databases and believed that which can improve the clinical application of detected isoforms

#### 2.2.2 Filtering of SNV/INDEL

There are also four steps to filter SNV/INDEL[Figure3].

The first step is to deplete the variant with sequencing depth below the confidence threshold. We calculate depth threshold as the formula below:

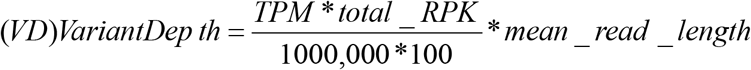

*RPK in the formula refers to Reads Per Kilobase, calculated by dividing the read counts by the length of each gene in kilobases*.

*TPM refers to Transcripts Per Kilobase Million,calculated by dividing the PRK values by the “per million” scaling factor*.

Considering that the peptide translated by variant transcripts with a TPM higher than 35 would be considered to have immunogenicity[20], we set the TPM to 35, and the threshold of Variant Depth can be calculated using the formula.

The second step is to remove background noise. For this purpose, we mainly use the column of FILTER in the VCF file to remove variants with flags such as P8, Q10, bias, etc.

The third step is to deplete variants derived from SNP or germline mutations by comprehensively analyzing the dbSNP annotation information and the allele frequency (AF). The last step is to annotate and identify mutations associated with clinical disease using a database such as COSMIC.

### 2.3 Benchmarking with biological reference standards

Initially, we wanted to benchmark all modules that detect novel variant isoform (NVI), but we only found a PacBio raw data of human brain reference spiked with 2% SIRV-Set 4, a kind of Biologics Reference Standards that can only be used to benchmark for ISOFORM calling. As an artificial isoform set, SIRV-set 4 contains 69 artificial transcript variants that mimic the splicing characteristics of 7 human model gene loci to comprehensively reflect variations of alternative splicing, alternative transcription start- and end-sites, overlapping genes, and antisense transcripts. SIRV-set 4 also contains 15 artificial transcript variants in five length categories (4 kb, 6 kb, 8 kb, 10 kb, and 12 kb, respectively) to mimic the length complexity of human transcripts.

The SIRV-set4 was developed to validate the performance of isoform-specific RNA-Seq workflows [27]. We download the original format data of Brain_Reference_SIRV_4_flnc.bam from the webpage (https://trace.ncbi.nlm.nih.gov/Traces/?view=run_browser&acc=SRR16762346&display=data-access). In the test run of the pipeline, we just provided the data path as input, without modifying any default parameters. All the analysis results can be harvested in about 4 hours in a high-performance computer cluster with Intel(R) Xeon(R) Platinum processor (8269CY CPU, 3.10GHz) and a Linux machine running Ubuntu 20.04.4 LTS. Figure 4 shows the distribution of time consumptions in different steps. Since the raw data of the sample is FLNC reads file that has been partially pre-processed, the quite time-consuming CCS step is omitted.

**Figure 4.**
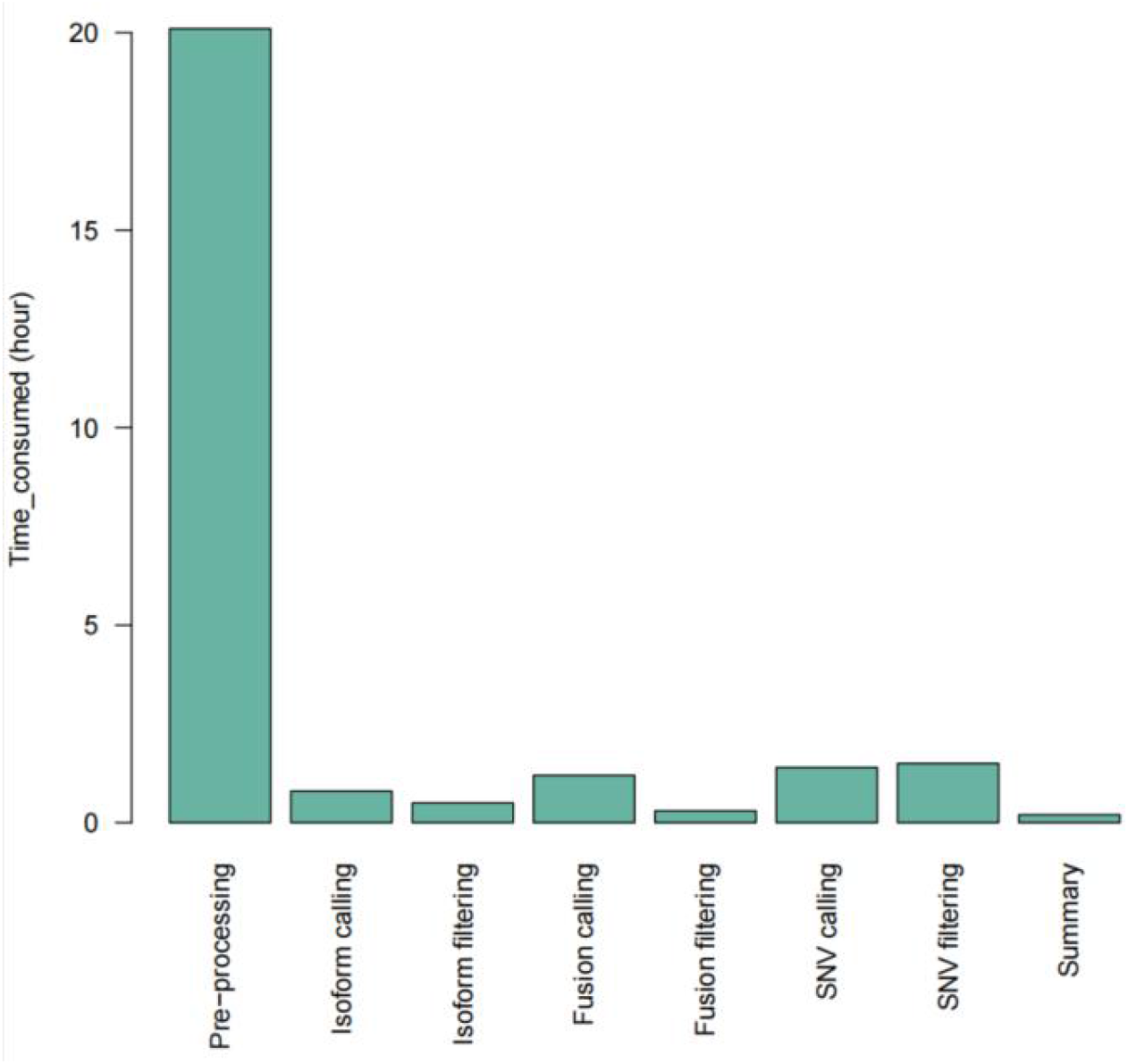
Time consumption per step.The pre-processing step of the sequences generated from 3^rd^ platform takes the longest time because of the alignment of the consensus sequence to obtaining the subreads from the polymerase reads. The time of other analysis steps is relatively short.

## 3 Results of biologics reference standards

The analysis of the SIRV-set4 took about 4 hours to complete. All modules were automatically analyzed and the corresponding analysis results were generated in their directories. Nevertheless, since the Biologics reference standards only have the information of the isoform (section 2.3), we only determined the accuracy of isoform calling module. We confirmed the calling results of all true isoforms listed in the contents table (https://www.lexogen.com/store/sirv-set4/) and calculated the recall ratio to determine whether the calling and hierarchical filtering steps of the analysis pipeline would erroneously miss the true positives. We found that we achieved a 100% (12/12) recall ratio (Table1,2) for the 12 long transcripts of SIRV-set4. However, for the detection of the 69 short SIRV transcripts, we only achieved a recall ratio of 89.9% (62/69), missing 7 isoforms.

**Table 1.** Short SIRV isoform of called and mis-called by SAKit.

| Short SIRV |  |  |  |  |
| --- | --- | --- | --- | --- |
| Transcript category ID | Total number of transcript | Recall number | Recall ratio | Missed |
| SIRV1 | 8 | 7 | 87.50% | SIRV105 |
| SIRV2 | 6 | 6 | 100.00% |  |
| SIRV3 | 11 | 10 | 90.90% | SIRV311 |
| SIRV4 | 7 | 6 | 85.70% | SIRV404 |
| SIRV5 | 12 | 10 | 83.30% | SIRV503,SIRV512 |
| SIRV6 | 18 | 17 | 94.40% | SIRV618 |
| SIRV7 | 7 | 6 | 85.70% | SIRV708 |
| Summary | 69 | 62 | 89.90% |  |

**Table 2.** Long SIRV isoform of called and mis-called by SAKit.

| Long SIRV |  |  |  |  |
| --- | --- | --- | --- | --- |
| Transcript category ID | Total number of transcript | Recall number | Recall ratio | Missed |
| SIRV10001 | 1 | 1 | 100.00% | / |
| SIRV10002 | 1 | 1 | 100.00% | / |
| SIRV10003 | 1 | 1 | 100.00% | / |
| SIRV12001 | 1 | 1 | 100.00% | / |
| SIRV12002 | 1 | 1 | 100.00% | / |
| SIRV12003 | 1 | 1 | 100.00% | / |
| SIRV4001 | 1 | 1 | 100.00% | / |
| SIRV4002 | 1 | 1 | 100.00% | / |
| SIRV4003 | 1 | 1 | 100.00% | / |
| SIRV6001 | 1 | 1 | 100.00% | / |
| SIRV6002 | 1 | 1 | 100.00% | / |
| SIRV6003 | 1 | 1 | 100.00% | / |
| SIRV8001 | 1 | 1 | 100.00% | / |
| SIRV8002 | 1 | 1 | 100.00% | / |
| SIRV8003 | 1 | 1 | 100.00% | / |
| Summary | 15 | 15 | 100.00% | / |

We investigated why those isoforms were missed by checking the results of each step, especially the preliminary results before calling or filtering.

- The 3^rd^ exon of the missed SIRV503 is too short: only 9bp; moreover, 2bp GT bases at the 5’ end were identical to the 2bp at the 5’ end of the 2^nd^ intron (Supplementary Figure 1A). Thus the last 7bp was misclassified as a softclip in alignment.
- The 1^st^ exon of the missed SIRV404 is also too short, only 31bp, and the 1^st^ intron is about 900bp (Supplementary Figure 1B), so the 1^st^ exon was misclassified as a softclip.
- SIRV708 was not detected because one of the introns was too short, only 20bp (Supplementary Figure 1C), so it was misclassified as deletion during alignment.

The root cause for all these missed isoforms is that the introns or exons are too short, resulting in a high mapping score in the global alignment to be misclassified as softclip or deletion. However, according to statistics of human RefSeq transcriptome, the overwhelming majority of exons and introns would be longer than 20bp, whereas the exons/intron of the 3 misclassified isoforms are 9bp, 31bp and 20bp, respectively--they could be regarded as extreme cases. The other four missing isoforms were not found in the raw data, indicating that they were lost in library construction or nucleotide sequencing.

All the results show that the SAKit pipeline has a satisfactory performance for identifying long isoform. In the presence of extremely short introns or exons, SAKit might miss some isoforms. Since the accuracy of isoform calling has a higher priority in practical application, we regard this issue as “acceptable residual risk”.

## 4 Conclusion

The concept of “neoantigen” has been prevalent in the new era of cancer immunotherapy. Neoantigens could be defined as immunogenic proteins/epitopes derived from aberrant genetic changes in tumor cells. Because normal cells do not have such aberrant genetic changes, neoantigens could be regarded as “absolutely” specific biomarkers of tumor cells. Therefore, in recent years biotech companies and research institutions have been actively developing immunotherapies against neoantigens. Nowadays, how to effectively identify neoantigens from clinical samples has become a valuable question to be addressed for translational research.

The “classical” approach for identifying neoantigens is based on searching nonsynonymous mutations in canonical open reading frames (ORFs). Such a “classical” approach is easily understandable, though its limitation is also apparent: only tumors with a high mutation burden have sufficient nonsynonymous mutations inside ORFs to induce novel proteins[28]; moreover, neoantigens with immunogenicity are only a tiny subset of novel proteins. Unfortunately, many cancer types have a low tumor mutation burden (<5 mutations/Mb), e.g., prostate cancer, glioblastoma, acute myeloid leukemia, etc. For these cancer types, it is barely possible to find sufficient neoantigens in canonical ORFs for immunotherapy [29].

However, due to aberrant genetic changes, most types of cancer cells involve alternative RNA splicings that do not exist in normal cells [30]. RNA alternative splicing provides another source of novel proteins, which are translated from the RNA sequences that are never translated in normal cells [28,31]. Therefore, identifying novel proteins and neoantigens from RNA alternative splicing would be an effective new approach to break the limitation of the “classical” approach, therefore greatly expand the application of immunotherapies against neoantigens.

As a powerful tool for studying peptidomes or proteomes, MS theoretically could identify novel proteins derived from RNA alternative splicing. In fact, the performance of MS heavily relies on the characteristics of reference databases. As discussed in the introduction section, the canonical human proteome is a disadvantageous reference database for identifying novel proteins that do not belong to canonical human proteome at all. To solve the logical paradox above, the prevalent HIFI and NGS technology provide a scientifically-sound method to establish the sample-specific reference database with hypothetical protein sequences derived from RNA translation.

Multiple mechanisms including fusion, SNP and INDEL, all contribute to the generation of novel RNA transcripts, as well as the subsequent induction of novel proteins. This fact calls for a powerful bioinformatic tool to process RNA sequencing data, and cover all types of RNA transcript variants. Therefore, we developed the SAKit pipeline that enables one-key running to comprehensively analyze the data of HIFI sequencing and NGS data. By far, the SAKit has the broadest coverage of all variants that can induce potential novel proteins than published workflows. And for the analysis results, we integrated and converted all types of variants into fasta-format protein sequences, which are very convenient for users to use for subsequent analysis, such as protein structure prediction, neoantigen immune epitope analysis, etc. In terms of resource usage, SAKit is capable of processing high-throughput data with limited computational power and time consumption. Moreover, its accuracy and reproducibility are ideal for processing clinical data. According to the benchmarking results with standard biological reference, the sensitivity of SAKit in long isoforms calling is as high as 100%, which shows that our SAKit fully utilizes the advantages of HIFI long-length reads sequencing. The detection sensitivity of short isoforms reaches 89.9%, which is lower than the detection sensitivity of long isoforms which seems unreasonable. However, among the 7 short isoforms that were missed, 3 of them have no raw reads supporting their structures in the original data at all, which means that those nucleotides of short isoforms may be degraded during the library construction. The other 4 isoforms are all due to the existence of extremely short introns or exons, resulting in almost no comparison during the alignment process. It is impossible to distinguish between intron/exon and deletion/insertion. These cases are all due to the highly artificial factors in the standard samples when they were designed, which are rare in the human transcriptome.

In conclusion, we have developed an all-in-one pipeline that can simultaneously detect all variants with high sensitivity and is user-friendly, thus paving the way for identifying novel proteins and neoantigens for immunotherapy.

---

**Supplementary Figure 1A.**
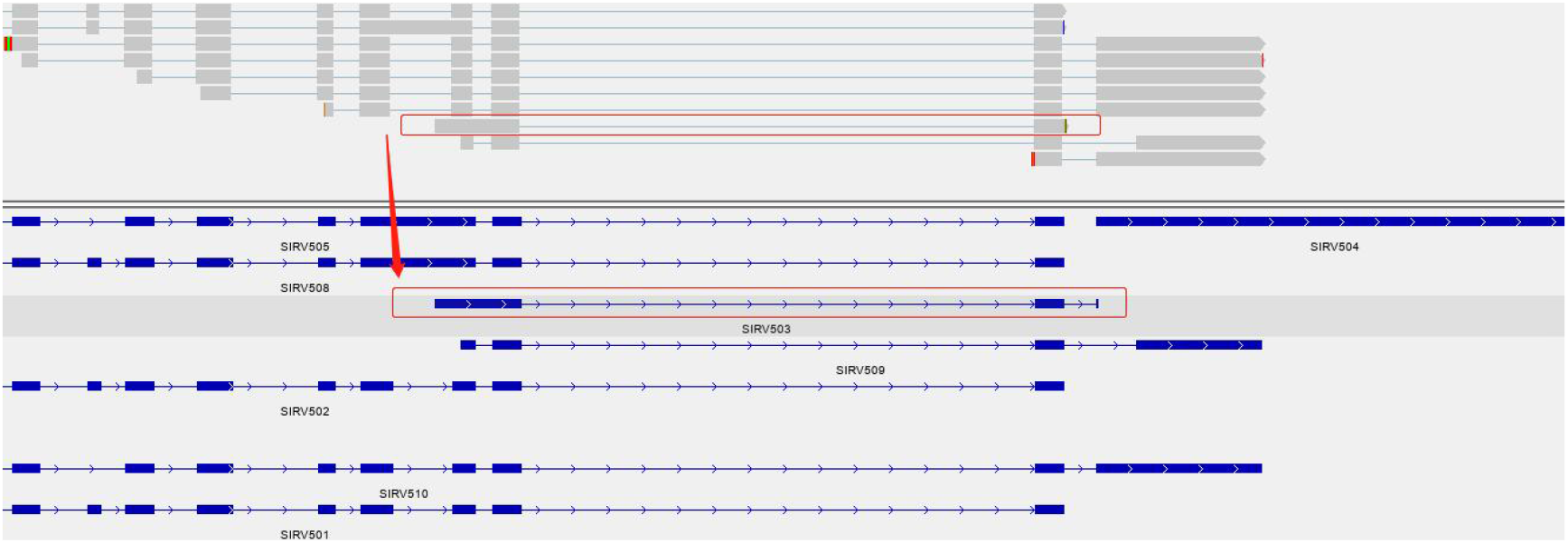
SIRV503

**Supplementary Figure 1B.**
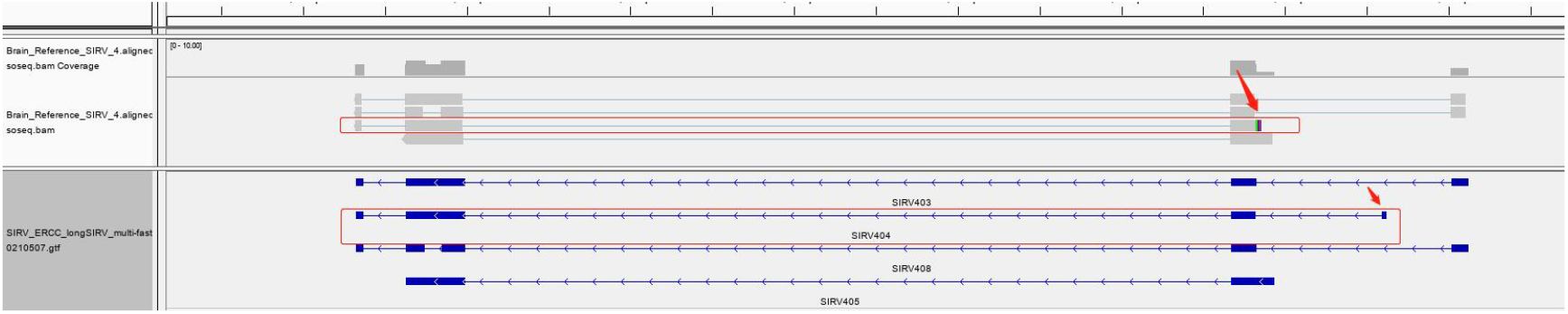
SIRV404

**Supplementary Figure 1C.**
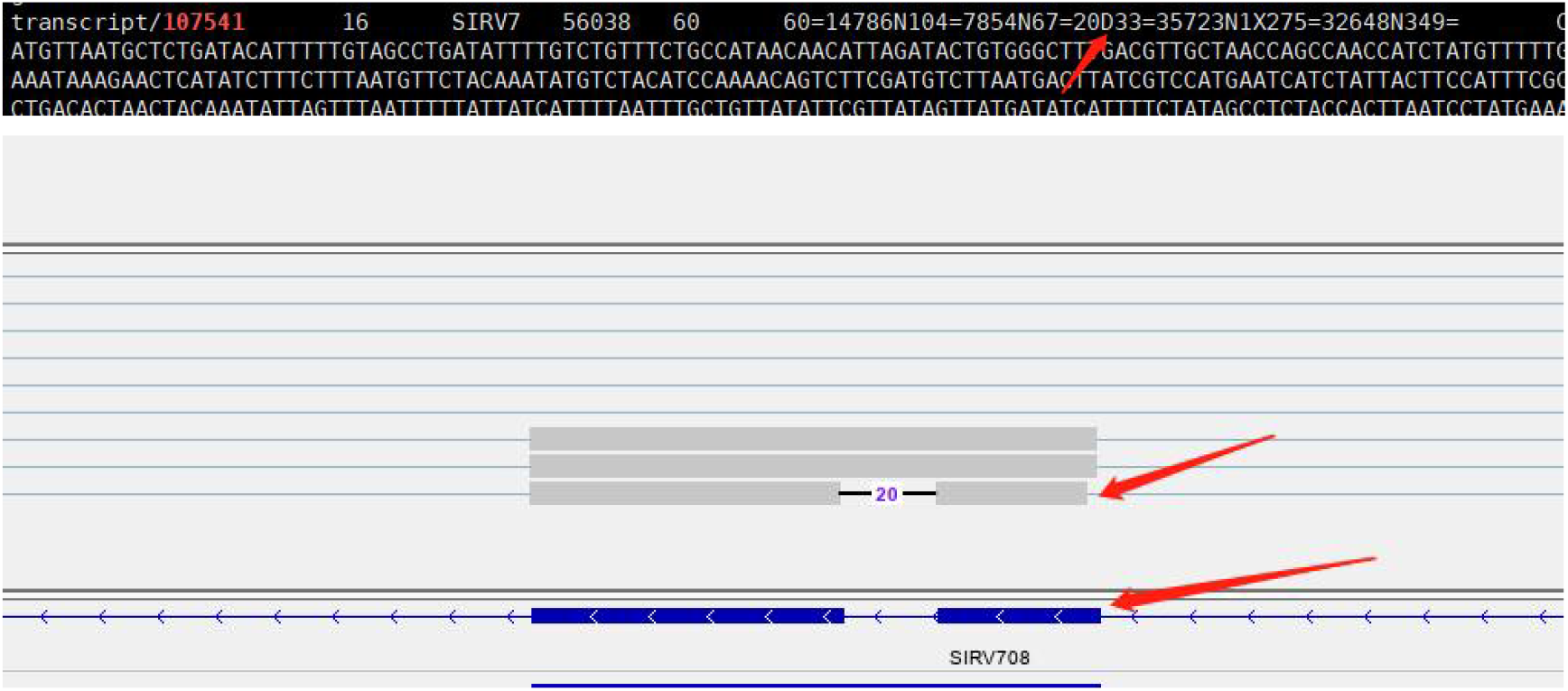
SIRV708

